# Lysosome-targeted lipid probes reveal sterol transporters NPC1 and LIMP-2 as sphingosine transporters

**DOI:** 10.1101/2021.11.10.468010

**Authors:** Janathan Altuzar, Judith Notbohm, Frank Stein, Per Haberkant, Saskia Heybrock, Jutta Worsch, Paul Saftig, Doris Höglinger

## Abstract

Lysosomes are central catabolic organelles involved in lipid homeostasis and their dysfunction is associated with pathologies ranging from lysosomal storage disorders to common neurodegenerative diseases. The mechanism of lipid efflux from lysosomes is well understood for cholesterol, while the export of other lipids, particularly sphingosine, is less well studied. To overcome this knowledge gap, we have developed functionalized sphingosine and cholesterol probes that allow us to follow their metabolism, protein interactions as well as their subcellular localization. These probes feature a modified cage group for lysosomal targeting and controlled release of the active lipids with high temporal precision. An additional photo-crosslinkable group allowed for the discovery of lysosomal interactors for both sphingosine and cholesterol. In this way, we found that two lysosomal cholesterol transporters, NPC1 and LIMP-2/SCARB2, also directly bind to sphingosine. In addition, we showed that absence of either protein leads to lysosomal sphingosine accumulation which suggests a sphingosine transport role of both proteins. Furthermore, artificial elevation of lysosomal sphingosine levels impaired cholesterol efflux, consistent with sphingosine and cholesterol sharing a common export mechanism.

## Introduction

Lipid homeostasis is maintained by an intracellular network of enzymes and regulated by organelle-to-organelle communication. Within this network, the lysosome plays a central role as host for degradation, recycling and trafficking of biomolecules such as proteins, nucleic acids and lipids (Ballabio & Bonifacino, 2020). Lysosomal dysfunction manifests in severe human pathologies known as lysosomal storage diseases (LSDs) which are often associated with lipid and particularly sphingolipid accumulation (Platt, 2018; Marques & Saftig, 2019).

Cholesterol is one well-studied lipid that relies on lysosomal trafficking. Dietary cholesterol reaches its target cells in plasma-low-density lipoprotein form (LDL). Upon endocytosis, LDL is hydrolysed in lysosomes, triggering release of free cholesterol (Brown & Goldstein, 1986). Subsequently, lysosomal cholesterol is transported to the plasma membrane (PM) or endoplasmic reticulum (ER). At the ER, sterol levels are sensed and cholesterol biosynthesis is regulated accordingly (Goldstein *et al*, 2006). Excess cholesterol is converted into cholesteryl esters for storage in lipid droplets (Thiele & Spandl, 2008). The exact mechanism of cholesterol egress from endocytic organelles to the ER or PM compartment is not fully elucidated. However, a substantial body of evidence implicates Niemann-Pick type C1 and C2 proteins (NPC1 and NPC2) in lysosomal cholesterol traffic (Xu *et al*, 2007; Infante *et al*, 2008; Nguyen Trinh *et al*, 2018; Meng *et al*, 2020). Aside from transporting cholesterol towards the lysosomal limiting membrane via a hydrophobic, intramolecular tunnel (Winkler *et al*, 2019). NPC1 has further been shown to form organelle contact sites with ER-resident proteins such as ORP5 (Du *et al*, 2011) or Gramd1b (Höglinger *et al*, 2019). This tethering function of NPC1 may further contribute to its importance in lysosomal cholesterol efflux from the lysosome to the ER. Alternatively, cholesterol can exit the lysosome through lysosomal integral membrane protein 2 (LIMP-2/SCARB2) which also features a hydrophobic intramolecular tunnel (Neculai *et al*, 2013).

While post-lysosomal cholesterol traffic has been extensively studied (Meng et al, 2020; Schoop et al, 2021), other lipids are also degraded in the lysosome. Here, the biologically active class of sphingolipids are of interest, given that a large number of LSDs feature lysosomal sphingolipid accumulation. Glycosphingolipid (GSL) catabolism proceeds through internalisation of the PM and the recycling of membrane components through the endo-lysosomal degradation pathway (Futerman & Riezman, 2005). Here, sphingomyelin (SM) and GSLs reach the lysosome and are degraded to form ceramides (Hannun & Obeid, 2008). In the last step of lysosomal sphingolipid catabolism, ceramides (Cer) are hydrolysed to form sphingosine (Sph) (Hannun & Bell, 1989), the backbone of all sphingolipids. Sphingosine can either be recycled back into the sphingolipid biosynthetic pathway (Kitatani et al, 2008) or phosphorylated to form sphingosine-1-phosphate and be degraded through the actions of sphingosine-1-phosphate lyase (SGPL1). Breakdown products of this degradation pathway can feed into the glycerolipid synthesis pathway (Bektas *et al*, 2010). While much is known about the enzymes involved in sphingolipid catabolism, the machinery responsible for lysosomal sphingolipid export is completely unknown. This can be attributed to a lack of functional tools to investigate sphingolipids on a single-organelle and single lipid species level.

To date, several tools such as fluorescent, isotope-labelled or bi-functional lipids have been used to follow sphingolipid metabolism and localisation (Haberkant *et al*, 2016; Bockelmann *et al*, 2018). Covalently-labelled fluorescent lipids contain fluorescent dyes such as fluorescein, nitrobenzoxadiazole (NBD), pyrene and BODIPY-like structures attached to their acyl chain or head group have been reported (Pagano *et al*, 2000; Schwarzmann *et al*, 2014). However, these constructs exhibit several drawbacks, such as aberrant membrane integration, mislocalisation, poor solubility and cell uptake (Ashcroft *et al*, 1980; Chattopadhyay, 1990; Maier *et al*, 2002). Bi-functional lipids (Haberkant *et al*, 2013, 2016) on the other hand have become popular compounds used to circumvent the problems associated with fluorescent lipids. These lipids combine a photocrosslinkable group, which allows crosslinking upon short-wavelength UV illumination, with an alkyne or azide group for subsequent staining or enrichment of lipid-protein complexes. One major drawback is that these lipids are rapidly metabolized after addition to cells, giving rise to a host of modified lipid metabolites. This hinders the study of small bioactive sphingolipids such as sphingosine, which, due to their potent signalling properties, are rapidly metabolized or degraded by the cell (Haberkant *et al*, 2016). To circumvent rapid metabolism, tri-functional lipids have been developed, where photocleavable protection (or ‘cage’) groups allow for equal loading of all cells before the active lipid probes can be released by a flash of longer-wavelength UV light (Höglinger *et al*, 2014, 2017). This strategy achieves a rapid burst of the lipid probe with high spatial and temporal resolution. Additionally, using an inherently fluorescent cage group such as coumarin allows the visualisation of the probe’s localisation before uncaging. However, it was shown that the cage group associates nonselectively with all internal membranes and therefore artificially mislocalizes the lipid of interest (Höglinger *et al*, 2017). As such, tri-functional lipids are not well-suited to study questions related to inter-organellar transfer. However, an additional strategy has recently been described to overcome this technical problem and achieves organelle specificity. Here, slight chemical modifications to the coumarin cage group led to their pre-localisation to different organelles such as the plasma membrane, mitochondria or lysosomes (Nadler *et al*, 2015; Wagner *et al*, 2018). So far, organelle-targeting of caged lipids has only been applied to non-functionalized or deuterated lipids (Wagner et al, 2018; Feng et al, 2019a, 2019b).

In this study, we combine the advantages of tri-functional and organelle-targeted lipids to create versatile tools for the visualisation of single lipids and for the identification of their protein interactome at the single-organelle level. We synthesised and applied lysosome-targeted photoactivatable and crosslinkable (pac) sphingosine (**lyso-pacSph**) and cholesterol (**lyso-pacChol**) to address outstanding questions regarding post-lysosomal sphingolipid trafficking and metabolism. These multi-functional tools allow us to (i) pre-localise sphingosine and cholesterol probes into lysosomes, (ii) screen for lysosomal protein-lipid interactors, (iii) follow their post-lysosomal metabolic fate by thin-layer chromatography (TLC) and (iv) visualise time-resolved lipid localization by confocal microscopy. We demonstrate the suitability of such a design by using lyso-pacSph to identify bona fide sphingosine transporters and show a surprising commonality in lysosomal sphingosine and cholesterol export routes.

## Results

### Synthesis and characterisation of lysosome-targeted pacSphingosine and pacCholesterol probes

We have designed lyso-pacSph and lyso-pacChol to feature four functionalities: (i) a photoactivatable diazirine ring for UV-dependent photocrosslinking, (ii) a clickable alkyne group which allows post-crosslinking functionalization of the lipid with fluorophores or biotin (Haberkant *et al*, 2013) and a (iii) photocleavable protecting group (cage) (Höglinger *et al*, 2017) equipped with a (iv) tertiary amine lysosomal targeting group. This targeting group is protonated at endo/lysosomal pH (4.5 – 6.5) and retained in the acidic environment of late endosomes and lysosomes (Kaufmann & Krise, 2007). This design ensures that all probe molecules are delivered to lysosomes before a flash of light (‘uncaging’) releases the active lipid species. We initially synthesised the lyso-coumarin caging group **(4)** from a previously described 7-ethylamino-coumarin precursor (Wagner *et al*, 2018) and added a dimethylamine-containing linker to incorporate the targeting group. Next, we coupled this cage to commercially available pacSph and pacChol (Hulce *et al*, 2013) through respective carbamate and carbonate linkages in order to afford the lyso-pacSph **(5)** and lyso-pacChol **(6)** with 99% and 96% yield, respectively **(Fig. 1A)**

**Figure 1.**
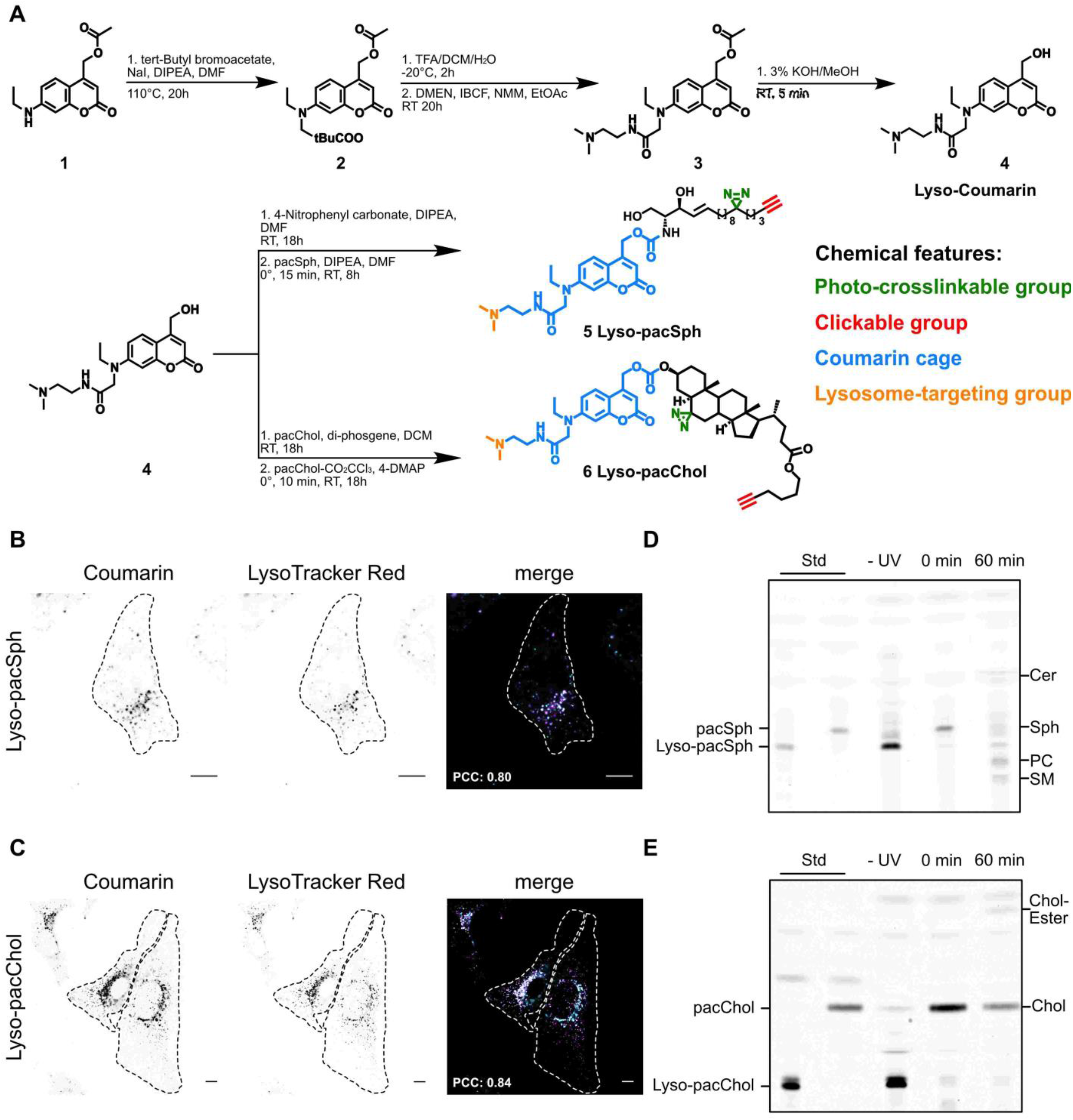
Synthesis, sub-cellular localisation and characterisation of lysosome-targeted pacSphingosine and pacCholesterol probes. **A.** Synthesis of lysosome-targeted coumarin cage (Lyso-Coumarin, **4**), lysosome-targeted pacSphingosine (Lyso-pacSph, **5**) and lysosome-targeted pacCholesterol (Lyso-pacChol, **6**) and annotation of the functional groups by color-coded legend. **B, C.** Lysosomal localization of lyso-probes. Confocal images of coumarin fluorescence in HeLa cells pulsed with Lyso-pacSph or Lyso-pacChol (10 μM) for 1h and 45 min respectively, chased overnight and incubated with LysoTracker Red (100 nM) 30 min previous to imaging. Scale bar: 10 μm. Pearson correlation coefficient (PCC) of non-thresholded images calculated using the Coloc 2 feature from ImageJ. Lyso-probe: cyan, LysoTracker Red: magenta. **D, E**. Thin-layer chromatography (TLC) of HeLa cells pulsed with Lyso-pacSph or Lyso-pacChol (10 μM) for 1h and 45 min respectively, chased overnight, harvested immediately and/or irradiated with a 405 nm UV light for 90s and further incubated at 37°C for indicated times. Cellular lipids were extracted, labeled with 3-azido-7-hydroxy coumarin by CuAAC click reaction, spotted and developed on a TLC silica gel plate.

Following synthesis of the compounds we investigated cellular uptake of these newly synthesised probes by live-cell confocal microscopy using the inherent fluorescence of the coumarin cage group. We found that both probes were readily taken up by HeLa cells over a wide range of concentrations from 750 nM to 10 μM after 60 mins of labelling **(Fig. EV1A)**. Interestingly, lyso-pacChol showed an immediate vesicular localization at all investigated concentrations, whereas lyso-pacSph was distributed to all internal membranes. To improve the subcellular localization of lyso-pacSph, we next varied probe-free incubation (‘chase’) times ranging from 0 min to 18 h **(Fig. EV1B)**. Lyso-pacChol remained located in vesicles at all chase times, but lyso-pacSph required longer chase times with exclusive vesicular localization obtained after 18 h of chase. Using these optimised, 18 h-chase conditions for both probes, we then investigated whether the vesicles stained with lyso-pacSph and lyso-pacChol were indeed lysosomes. We found that the fluorescent coumarin pattern of the lyso-probes overlapped completely with LysoTracker™ signal as quantified using Pearson’s correlation coefficient (PCC) with values of 0.84 or higher **(Fig. 1B and C)**.

Given that long chase times were required to achieve optimal lysosomal localisation of lyso-pacSph, we next investigated the stability of the lyso-probes to withstand the activities of lipases and esterases found in the lumen of lysosomes prior to their uncaging. To this end, we pulsed and chased HeLa cells with both lyso-probes for up to 24 h, extracted the lipids and added a commercially available fluorophore to the alkyne group by means of click chemistry. TLC analysis showed that no additional bands appeared during 24 h of chase times **(Fig. EV1C and D)**, confirming that the lyso-probes are inert and not subject to cellular metabolism whilst in their caged form. To evaluate the photo-cleavage efficiency of the lyso-probes, we performed uncaging experiments of both lyso-probes in aqueous solution with increasing duration of UV irradiation (405 nm). Here, TLC analysis showed that 60-90 seconds were sufficient to photocleave (‘uncage’) almost all probe molecules and release the active lipid species **(EV1E)**. Having optimized the uncaging step, we next studied the metabolic consequences of uncaging the lipid probes in living cells. TLC analysis of lyso-pacSph **(Fig. 1D)** and lyso-pacChol **(Fig. 1E)** again showed that incubation without uncaging (-UV) does not release the active (pac) lipid species, whereas immediately after uncaging (0’) only the active species pacSph and pacChol could be detected. When incubating a further 60 min after uncaging (60’) however, the modified lipids were further incorporated into their respective downstream metabolites. Lyso-pacSph was converted to ceramide (Cer), sphingomyelin (SM) and to a significant degree to phosphatidylcholine (PC), a product of the sphingosine-1-phosphate lyase (S1PL)-dependent breakdown pathway (for identification of the bands with their respective standards, see **Fig. EV1F**). In further experiments, to avoid the incorporation of the labelled sphingosine backbone into abundant glycerolipids, a SGPL1-deficient HeLa cell line (Gerl *et al*, 2016) was employed, thereby limiting experimental readouts to lipid species containing the sphingosine backbone. Lyso-pacChol on the other hand was metabolized fairly slowly and to only one further species, a cholesterol-ester. Therefore, all further experiments using Lyso-pacChol were carried out in HeLa WT cells.

Together, these data showed that the sphingosine and cholesterol analogues released inside lysosomes closely resemble their endogenous counterparts and readily participate in their respective metabolic pathways, in accordance with previous studies using tri-functional sphingosine (Höglinger *et al*, 2017) and bi-functional cholesterol (Höglinger *et al*, 2019).

### Suitability of Lyso-pacSph and Lyso-pacChol to investigate lysosomal protein-lipid interactions

We next investigated the usefulness of these stable lysosome-targeted probes in studying subcellular protein-lipid interactions. To this end, we compared the non-caged pacSph and pacChol probes (Haberkant *et al*, 2016; Hulce *et al*, 2013), which upon addition to the medium are taken up and globally distributed throughout the cell, with our newly synthesised lyso-probes. Both versions of probes were subjected to a workflow consisting of metabolic labelling, uncaging and crosslinking photoreactions, enrichment of the protein-lipid complexes by streptavidin-mediated immunoprecipitation and analysis by Western Blot **(Fig. 2A)**. In this way, we could reveal all biotinylated protein-lipid complexes by using fluorescent streptavidin **(Fig. 2B)**. As expected, the lysosomal probes gave rise to a reduced subset of cross-linked partners compared to their globally distributed counterparts. However, several bands (marked with red arrows) could only be detected using the lysosomal probes, highlighting the sensitivity of this method to capture scarce or brief interactions which would be missed when using non-prelocalised probes.

**Figure 2.**
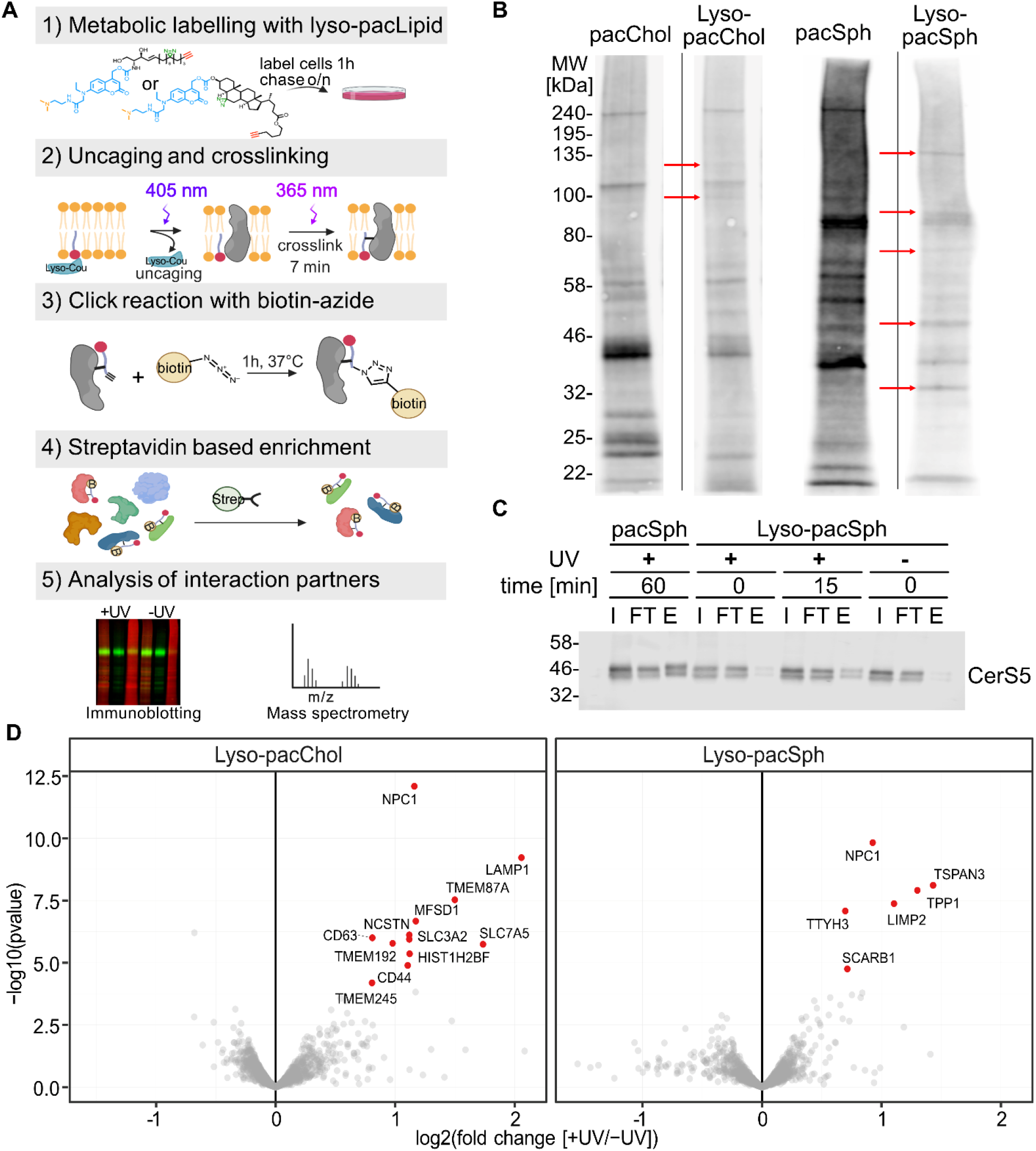
Application of lyso-pacChol and lyso-pacSph for identification of protein-lipid interactions. **A.** Graphical illustration of workflow using lysosome-targeted probes to capture protein-lipid interactions. **B.** Differential identification of proteins crosslinked to cholesterol probes and sphingosine probes. Biotinylated protein-lipid complexes were visualized using fluorescently-labelled Streptavidin. Proteins identified with lyso-probes but not with globally distributed probes are marked with red arrows. **C.** Time-dependent identification of FLAG-tagged ceramide synthase 5 (CerS5) by lyso-pacSph compared to globally distributed pacSph. HeLa SGPL1-/- cells were labelled with lyso-pacSph (5μM) or pacSph (2μM, 1h), uncaged and either crosslinked immediately or incubated 15 mins before crosslinking and subjected to the workflow in (A) featuring crosslinking (+UV) or no crosslinking (-UV) steps. The inputs (I, 10%), flow-throughs (FT, 10%)and eluates (E, 100%) were immunoblotted against FLAG-tag. **D.** Chemoproteomic analysis of lyso-pacChol and lyso-pacSph interactors. Volcano plot showing the results of a differential abundance analysis using imma package (moderated t-test, p-values estimated by fdrtool package) of proteins crosslinked to the respective probes. Log2-fold change of crosslinked over non-crosslinked (x-axis) and negative log10 P-values (y-axis) of protein interactors. Hit-proteins (red annotated dots) displayed a false-disovery rate of ≤ 0.01 and a fold-change of > 1.5

Next, we tested whether this workflow was suitable to enrich known lysosomal proteins and investigated the well-studied lysosomal cholesterol-binding proteins NPC1, LAMP1 and LIMP-2/SCARB2 by immunoblotting following Streptavidin-mediated immunoprecipitation **(Fig. EV2A).** Encouragingly, all three investigated proteins could be detected in eluate (‘E’) fractions of both pacChol as well as lyso-pacChol treated samples, whereas they were absent in the eluate fractions of –UV samples, where the photocrosslinking step was omitted. This supports that the established workflow can specifically detect lysosomal protein-lipid interactions.

As a further challenge to the spatio-temporal resolution, we next studied the availability of lyso-probes to non-lysosomal proteins. To this end, we investigated whether lyso-pacSph was able to interact with an ER-resident protein, ceramide synthase 5 (CerS5, **Fig. 2C**). Given that sphingosine is a substrate of CerS5, we unsurprisingly detected CerS5 in the eluate fractions of samples treated with the globally distributed pacSph. However, samples collected immediately upon uncaging of lyso-pacSph (0 min chase) did not reveal CerS5 in the elution fraction, further supporting the exquisite time-sensitivity of this assay. However, when a 15 min incubation was added upon uncaging, CerS5 could once again be detected in the elution fraction, indicating a successful transport of the lyso-pacSph probe to the ER. Together, these data highlight the suitability of the lyso-probes to specifically detect lipid interacting proteins with a high spatial and temporal control.

### Chemoproteomic profiling of Lyso-pacSph and Lyso-pacChol

Next, we set out to explore lysosomal interactors of sphingosine and cholesterol in an unbiased fashion. At 0 min post-uncaging to ensure maximum lysosomal localization, we again used the lyso-probes in the workflow based on streptavidin-mediated immunoprecipitation, but now we analysed the eluates by tandem-mass tag (TMT)-based quantitative proteomics. Comparison of crosslinked (+UV) with non-irradiated (-UV) samples gave a small subset of proteins which passed the strict threshold of a false discovery rate (FDR) below 0.01 and a fold-change of 1.5 **(Fig. 2D,** for a list of hit proteins, see **Fig. EV2C and D)**. For lyso-pacChol-treated cells, we found known lysosomal cholesterol interactors such as NPC1, LAMP1 (Kwon *et al*, 2009; Li & Pfeffer, 2016) among 12 candidate hits. Only six of those had been previously identified using the non-prelocalized pacChol probe (Hulce *et al*, 2013). Lyso-pacSph-treated samples gave only six candidate proteins. Two of those proteins (SCARB1 and TTYH3) are annotated as plasma membrane proteins. Among the other four candidates, we found NPC1 and LIMP-2/SCARB2 as potential sphingosine interactors, two proteins that were previously implicated in lysosomal cholesterol egress. We next validated the sphingosine interaction with NPC1 and LIMP-2/SCARB2 by immunoprecipitation and Western Blot analysis and indeed found that both probes were efficiently pulled down with both, the globally distributed pacSph as well as with lyso-pacSph **(Fig. EV2B)**, further strengthening both proteins as novel sphingosine-interactors.

### Metabolic fate of lysosomal cholesterol and sphingosine

Next, we were curious to study the metabolic fate of the lyso-probes and, in particular, whether the newly identified sphingosine interactors NPC1 and LIMP-2/SCARB2 affected the kinetics of post-lysosomal cholesterol and sphingosine metabolism. To this end, we performed pulse-chase experiments in which we incubated WT, NPC1^−/−^ (Tharkeshwar *et al*, 2017)and LIMP-2/SCARB2^−/−^ HeLa cells with both lyso-probes for different times after uncaging and analysed the resulting modified lipids by thin-layer chromatography. Of note, this experiment does not feature a crosslinking step and therefore the time-resolution is decreased as additional minutes are needed to collect cells and perform lipid extraction. As expected, lyso-pacChol-labelled cells showed a time dependent increase in cholesterol-ester **(Fig. 3A)**. This progressive incorporation of cholesterol into cholesterol-esters is severely reduced and delayed in NPC1^−/−^ as well as LIMP-2/SCARB2^−/−^ HeLa cells (**Fig. EV3A and B** quantified in **Fig. 3B**) in line with previous studies showing a lysosomal export defect of cholesterol under these conditions (Hölttä-Vuori et al, 2008; Heybrock et al, 2019). WT Cells labelled with lyso-pacSph showed extremely rapid metabolic conversions **(Fig. 3C)** where the precursor was incorporated into higher sphingolipids such as ceramide, but also degraded via the S1PL-pathway, yielding a small portion of labelled phosphatidylcholine and phosphatidylethanolamine at the earliest timepoint. Conversely, NPC1^−/−^ cells showed less rapid conversions at the earliest timepoint as shown by high initial sphingosine levels in the quantification of sphingosine as percentage of total labelled lipids **(Fig. 3D and Fig. EV3C)**. Furthermore, NPC1^−/−^ cells exhibited a delayed decrease in Sph levels compared to WT cells, consistent with a potential transport defect. LIMP-2-deficient cells, on the other hand, featured slightly elevated initial sphingosine levels (**Fig. EV3D**) but their time-dependent decrease followed the same kinetics as in WT cells. Together, loss in lysosomal proteins NPC1 and LIMP-2/SCARB2 significantly impacted post-lysosomal cholesterol metabolism, whereas sphingosine metabolism was found to generally proceed much faster and depend more on NPC1 than LIMP-2/SCARB2.

**Figure 3.**
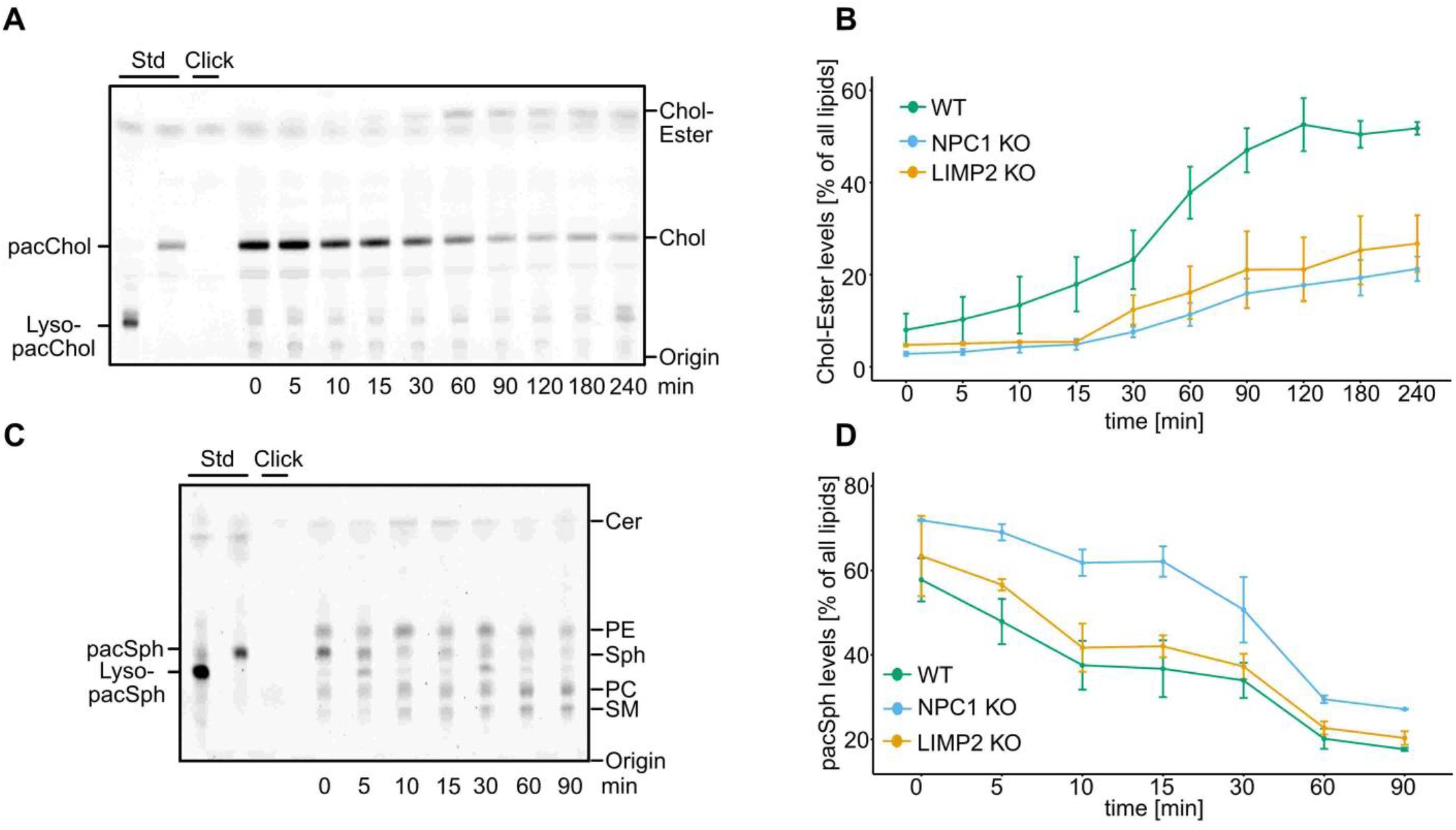
Lyso-pacChol and Lyso-pacSph metabolism in WT, NPC1 KO and LIMP2 KO cells by thin-layer chromatography. **A, C.** Post lysosomal metabolism of Lyso-pacChol **(A)** and Lyso-pacSph **(C)**. HeLa WT cells were labelled with Lyso-pacChol or Lyso-pacSph (10 μM) for 45 and 1h respectively and chased overnight. Upon uncaging, cells were chased and lipids extracted at indicated times, clicked with 3-azido-7-hydroxycoumarin and visualised by thin-layer chromatography. **B,D.** Quantification of post lysosomal cholesterol metabolism and sphingosine export. **B.** Quantification of cholesterol esterification in WT, NPC1 and LIMP2 -deficient HeLa cells. The intensity of the bands corresponding to pacChol and pacChol-Ester were added for each timepoint and the intensity of the ester-band with respect to the sum of both was displayed as percentage. **C.** Quantification of lysosomal sphingosine (Sph) export in WT, NPC1 and LIMP2-deficient HeLa cells. Sph is readily metabolized to ceramide (Cer) and sphingomyelin (SM) via the biosynthetic sphingolipid pathway, but also top phosphatidylethanolamine (PE) and phosphatidylcholine (PC) via the SGPL1-breakdown pathway. Intensity of the Sph is expressed as percentage compared to the sum of all labelled lipids. All values were calculated for each timepoint in three independent experiments. Data are shown as mean ± standard error.

### Lysosomal egress and subcellular localisation of sphingosine and cholesterol across organelles in NPC1 and LIMP-2/SCARB2 deficient cells

Next, we visualised the time-dependent changes in subcellular localisation of lyso-probes by confocal microscopy in order to investigate whether lysosomal egress was indeed delayed as suggested by the TLC studies. To this end, we took advantage of the versatility of the clickable-group to attach a fluorophore as another reporter molecule in pulse-chase experiments **(Fig. 4A and C)**. In WT cells labelled with lyso-pacChol, a predominantly lysosomal localisation was visible at 0 mins, followed by staining of internal membranes, the Golgi apparatus and the plasma membrane after 30 mins. At 60-90 mins, some of the probe localised to lipid droplets (for co-localisation with respective organelle markers, see **Fig. EV4.1A**), likely in form of cholesterol esters. In accordance with previous studies (Carstea, 1996; Saha *et al*, 2020), cells lacking NPC1 showed a significant retention of the lyso-pacChol probe in lysosomes for up 90 mins **(Fig. 4B)**. LIMP-2/SCARB2^−/−^ cells on the other hand showed an intermediate phenotype with internal membranes stained with similar kinetics as in WT cells, yet some lysosomes were visible throughout the 90 mins time course (**Fig. EV4.1C**, quantification in **Fig. 4B**). This could potentially point towards different populations of lysosomes, one population with a functional cholesterol export route (arguably via NPC1 to the ER) and a smaller population with defective cholesterol export mediated by LIMP-2/SCARB2. We next utilised lyso-pacSph to study the yet unexplored lysosomal export of sphingosine (**Fig. 4C**, for co-localisation see **Fig. EV4.2**). In WT cells, sphingosine re-localised from lysosomes to predominantly Golgi membranes at 30 mins (**Fig. EV4.2A and B**) while at later timepoints, much of the initial signal intensity was lost, consistent with efflux or effective catabolism of the sphingosine probe. In NPC1^−/−^ cells, a striking sphingosine storage phenotype was observed with lysosomes stained for the entire time course and only a small part of the probe localizing to other internal membranes (**Fig. EV4.2C**). This observation was corroborated by a continuously high Pearson coefficient throughout the time course **(Fig. 4D)**. LIMP-2/SCARB2^−/−^ cells, again, showed an intermediate phenotype with a partial export to internal membranes and partial storage in a small subset of lysosomes (**Fig. EV4.2D**). Together, these data show another surprising commonality between sphingolipid and cholesterol trafficking such that absence of previously known cholesterol transporters NPC1 and LIMP-2/SCARB2 also affect the kinetics of sphingosine export.

**Figure 4.**
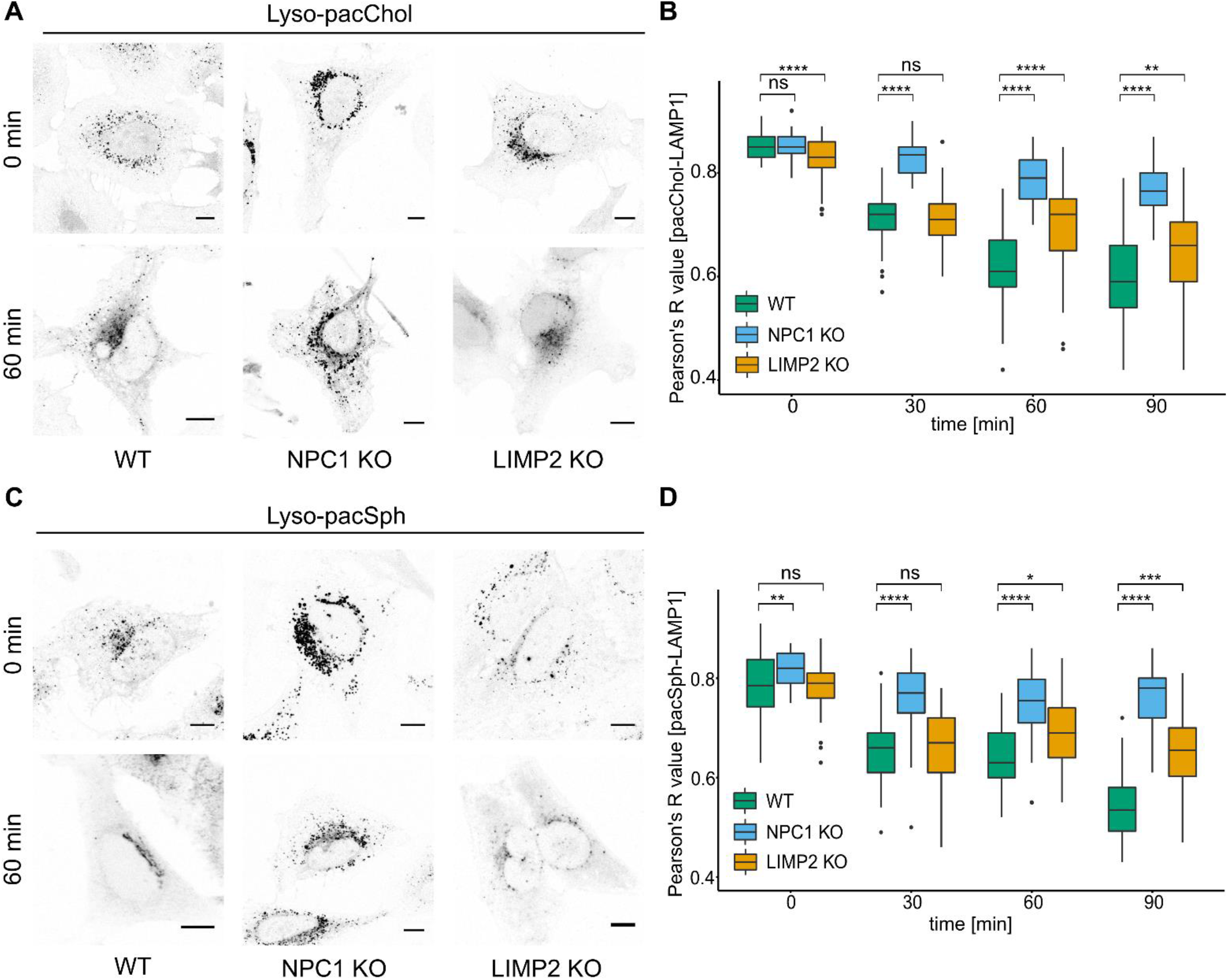
Visualization of lysosomal lipid egress in WT, NPC1 KO and LIMP2 KO cells. Subcellular localization of Lyso-pacChol **(A)** and Lyso-pacSph **(C)** in WT, NPC1 and LIMP2-deficient HeLa cells. Confocal images of cells labeled with Lyso-pacChol or Lyso-pacSph (10 μM) for 45 min and 1h respectively and chased overnight. Upon uncaging, cells were chased for the indicated times, crosslinked, fixed and functionalised with AlexaFluor 594-Picolyl-Azide **(A)** or AlexaFluor 488-Picolyl-Azide **(C)**. Scale bar: 10 μm. **B, D**. Quantification of lysosomal lipid egress. Pearson’s R value of non-thresholded images from lipid channel vs. LAMP1 immunofluorescence, calculated for each timepoint (n ≥ 45) using the Coloc 2 feature from Fiji. (ns P-value > 0.5 *P ≤ 0.05 **P-value ≤ 0.01 ***P-value ≤ 0.001 ****P-value ≤ 0.0001)

### Sphingosine transport defect is a direct consequence of NPC1 dysfunction

To further investigate the prominent sphingosine storage phenotype in NPC1-deficient cells and to exclude that the observed phenomenon was a downstream effect caused by long-term adaptation in our particular NPC1^−/−^ cell line, we next evaluated pharmacological inhibition of NPC1 in WT cells by acute treatment with U18666A. This drug has been shown to inhibit the sterol-sensing domain of NPC1 and to block cholesterol export (Lu *et al*, 2015). Repeating the lyso-pacSph pulse-chase experiment in the presence of U18666A showed that, again, sphingosine export from lysosomes was severely delayed compared to WT cells (**Fig.5A**, quantified in **Fig. 5B**), albeit to a slightly lesser degree than in a genetic knock-out model. To challenge whether the observed sphingosine storage was a primary effect due to loss of NPC1 or a secondary effect created by NPC1-induced cholesterol accumulation and a potentially resulting reduced capacity of lysosomes to export sphingosine, we subjected NPC1^−/−^ cells to starvation by growing them in lipoprotein-deficient medium. This treatment drastically reduced lysosomal cholesterol levels as revealed by Filipin staining **(Fig. EV5A**, quantified in **Fig. EV5B)**. Investigating lyso-pacSph efflux under these conditions revealed that sphingosine still remained trapped in lysosomes for the duration of the time course in the same degree as non-starved NPC1 cells **(Fig. 5A, quantified in 5B)**, arguing for a direct effect of NPC1 on sphingosine transport. To further challenge this hypothesis, we attempted to recreate an NPC1-like phenotype in a WT background by artificially increasing lysosomal lipid levels. To this end, we synthesised new lysosomally pre-localised probes lyso-Sph and lyso-Chol **(Fig. 5C)**. These probes feature the same cage group and lysosomal targeting groups, but lack the crosslinkable and clickable groups found in lyso-pacSph and lyso-pacChol. Applying these probes in combination with lyso-pacSph and lyso-pacChol, respectively, allowed us to pre-load lysosomes with two different lipids at the same time. Upon uncaging, both lipid types were released at the same time, yet we followed only the localisation of the lyso-pacLipids by virtue of their clickable group (schematically illustrated in **Fig. 5D**). Initially we applied lyso-pacSph together with a 20-fold excess of lyso-Chol to investigate whether sphingosine export was delayed in presence of a large excess of cholesterol **(Fig. 5E)**. Co-localisation analysis between the sphingosine probe and LAMP1 as performed by Pearson’s correlation coefficient **(Fig. 5F)** showed no detectable difference in sphingosine export with or without excess cholesterol. When inverting the experiment, i.e., flooding the lysosome with a 20-fold excess of sphingosine and following the export of lyso-pacChol, however, we could detect a delay in cholesterol export in the presence of sphingosine (**Fig. 5G**, quantified in **Fig. 5H**). These data further support the hypothesis that the sphingosine storage phenotype observed in NPC1^−/−^ cells was not a secondary consequence of cholesterol accumulation, but rather that increasing lysosomal sphingosine concentration induced an NPC1-like delay in cholesterol export.

**Figure 5.**
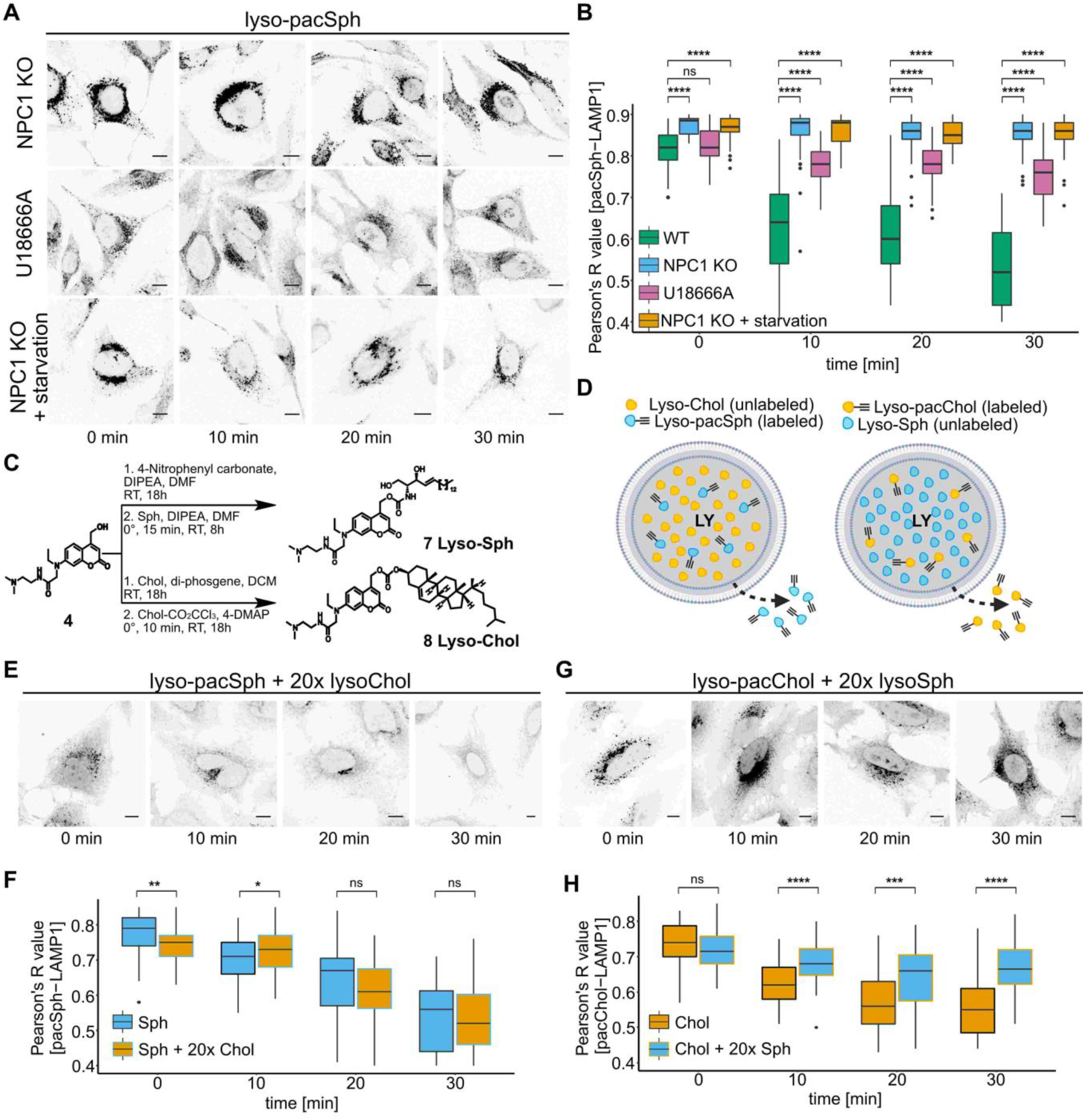
Sphingosine accumulation is a direct effect from loss of NPC1 and can re-create an NPC1-like phenotype. **A.** Confocal images of Lyso-pacSph in cellular models of NPC1. HeLa NPC1 KO cells (NPC1), HeLa WT cells treated with 0.5 μg/mL U18666A 24h prior to imaging (U18666A) and HeLa NPC1 KO cells incubated with lipoprotein deficient medium 48h prior to imaging (Starvation) were labeled with Lyso-pacSph (1.25 μM) for 1h and chased overnight. Upon uncaging, cells were chased from 0 to 30 min, crosslinked, fixed with methanol and functionalised with AlexaFluor 594-Picolyl-Azide. Scale bar: 10 μm. **B**.Quantification of lysosomal lipid egress. Pearson’s R value of non-thresholded images from lipid channel vs LAMP1 immunofluorescence, calculated for each timepoint (n ≥ 42) using the Coloc 2 feature from Fiji. (ns P-value > 0.05 *P ≤ 0.05 **P-value ≤ 0.01 ***P-value ≤ 0.001 ****P-value ≤ 0.0001). **C.** Synthesis of lysosome-targeted Sphingosine (Lyso-Sph, **7**) and lysosome-targeted Cholesterol (Lyso-Chol, **8**). **D** Recreation of an NPC1-like phenotype. Illustrative model of the performed experiment. **E,G.** Confocal images from HeLa WT cells labelled with either Lyso-pacSph (1.25 μM) and Lyso-Chol (25 μM) **(E)** or Lyso-pacChol (750 nM) and Lyso-Sph (15 μM) **(G)** for 1h and chased overnight. Upon uncaging, chase-crosslinking experiments were performed from 0 to 30 min. Cells were fixed with methanol and functionalised with AlexaFluor 594-Picolyl-Azide **(E)** or AlexaFluor 488-Azide **(G)**. Scale bar: 10 μm. **F,H.** Quantification of lysosomal lipid egress in **E** and **G**. Pearson’s R value of non-thresholded images from lipid channel vs LAMP1 immunofluorescence, calculated for each timepoint (n ≥ 42) using the Coloc 2 feature from Fiji. (ns P-value > 0.05 *P ≤ 0.05 **P-value ≤ 0.01 ***P-value ≤ 0.001 ****P-value ≤ 0.0001).

## Discussion

In this study, we combine the concepts of tri-functional lipids with organelle targeting to achieve exquisite spatiotemporal control over these probes. Upon pre-localisation of lipids within lysosomes in their caged and therefore inactive state, rapid release by a flash of light provides a synchronized starting point for tightly controlled pulse-chase experiments. This is particularly important for the study of signalling active lipids which are subject to rapid metabolism and degradation. In principle, this design is not restricted to lysosomes. The chemistry required for targeting other organelles such as the plasma membrane, ER, lipid droplets, mitochondria and Golgi is well established (Xu *et al*, 2016) and has even been applied to the pre-localisation of lipids (Wagner *et al*, 2018; Feng *et al*, 2019a). The tremendous stability of pre-localised lipids to withstand hydrolytic enzymes in their caged state - up to 24 h in this study – allowed the optimization of chase times to achieve exclusive organellar localisation. To our surprise, the small and soluble lipid lyso-pacSph required long chase times (> 4 h, best localisation at 18 h) for optimal lysosomal staining whereas the bulkier lyso-pacChol probe already displayed lysosomal localization after short labelling times. The reasons for such differential targeting kinetics are unclear. The lysosomotropism of exogenous molecules is dependent on a number of physicochemical parameters (such as pKa and lipophilicity) such that the behaviour of novel compounds is difficult to predict. However, this is a large area of research, especially in pharmacokinetics, where detailed mathematical models exist to explain these phenomena (Zhang *et al*, 2008).

Having optimized the pre-uncaging localizations of all probes, we applied our lyso-pac-lipids to studying the mechanisms of sphingosine exit from lysosomes. This is an important step in sphingolipid homeostasis and it is estimated that the so-called salvage pathway, that is, the reutilization of sphingosine in sphingolipid biosynthesis, utilises up to 50-90 % of the lysosomally-derived sphingoid base pool (Gillard et al, 1998; Kitatani et al, 2008). However, to date to the best of our knowledge, the molecular mechanism of lysosome-ER traffic of spingosine for both the salvage but also the SGPL1-dependent breakdown pathway is completely unclear. A recent study in yeast identified an ER-lysosomal tethering protein, Mdm1 which facilitated the incorporation of lysosomal sphinganine into dihydroceramide (Girik *et al*, 2021). This points towards a requirement for organelle contact sites in the traffic of Sph and argues for the presence of dedicated sphingosine transporters, similar to other lipids such as cholesterol (Antonny *et al*, 2018) or diacylglycerols (Saheki *et al*, 2016) which depend on the actions of lipid transport proteins such as OSBP or E-Syts for their trafficking between organelles.

To our surprise, our chemoproteomic screen identified two well-known cholesterol-binding proteins, NPC1 and LIMP-2/SCARB2, as sphingosine interactors. Both proteins have been described to feature hydrophobic cavities (‘tunnels’) through which cholesterol can pass (Neculai et al, 2013; Winkler et al, 2019). Interestingly, LIMP-2 was found to bind other lipids besides cholesterol such as PS (Conrad *et al*, 2017), hinting a broader specificity. NPC1 on the other hand, has not yet been studied with respect to its substrate specificity. However, patients suffering from NPC1 disease feature accumulation of not just cholesterol, but multiple lipids within lysosomes. Most notably, the levels of Sph are significantly increased in multiple tissues in a manner that is unique to NPC1 disease compared to all other sphingolipidoses (Rodriguez-Lafrasse *et al*, 1994). Together with our finding that lyso-pacSph crosslinked to NPC1, it seems likely that the lipid binding specificity of NPC1 goes beyond sterols as well. Based on the structures of both proteins, NPC1 (Winkler *et al*, 2019) and LIMP-2/SCARB2 (Neculai *et al*, 2013), we hypothesize that sphingosine could fit into the respective cholesterol binding pocket, which raises exciting questions about how this competition and the relative levels of the two lipids in the lysosome would influence their mechanism of transport.

We then investigated whether NPC1 could not only bind, but also traffic Sph out of the lysosomes. The notion of NPC1 as a Sph transporter was proposed following the findings that exogenous addition of Sph could induce an NPC1 phenotype in healthy cells and that pharmacological inhibition of NPC1 in cell culture resulted in sequential elevation of multiple lipids with Sph being the first to accumulate (Lloyd-Evans *et al*, 2008). However, other studies featuring pulse-chase experiments with radiolabelled sphingosine did not find evidence of a Sph trafficking defect in NPC1-deficient cell models (Blom *et al*, 2012), potentially because of the hour-long time courses employed. In our hands, several lines of investigations support the hypothesis that NPC1 is capable of trafficking Sph. First, metabolic labelling studies in NPC1^−/−^ showed a delayed post-lysosomal metabolism of Sph. Secondly, visualization of lyso-pacSph revealed a strikingly prolonged lysosomal staining in NPC1^−/−^ or U18666A-treated cells upon uncaging. In fact, Sph was stained within lysosomes for the entire duration of the experiments (going up to 4h) while WT cells cleared all lysosomal lyso-pacSph within 30 mins upon uncaging. And thirdly, we could confirm that uncaging of an excess of lysosomally-prelocalized Sph indeed led to delayed export of cholesterol, whereas the inverse experiment (releasing excess cholesterol) did not cause any delay in Sph export. This last finding could suggest that Sph is preferentially trafficked by NPC1 or that the binding of Sph to NPC1 is stronger than that of cholesterol. It is tempting to speculate about a model in which NPC1 is a cholesterol-regulated protein that is able to move other substrates, including sphingosine, to the limiting membrane of the lysosome for transport at lysosome-ER contact sites(Höglinger *et al*, 2019)

In conclusion, the presented method to study single lipids with subcellular resolution gave new insights into the previously understudied lipid Sph. Similar to cholesterol, Sph can exit the lysosomes via at least two proteins LIMP-2/SCARB2 and NPC1, likely using their hydrophobic tunnels for delivery to the limiting membrane of the lysosome. From there, it is tempting to speculate that dedicated lipid transport proteins would also shuttle Sph from lysosomes to other organelles at dedicated contact sites. The current dataset did not give hints towards potential inter-organellar transporters, potentially because the crosslinking was performed immediately upon uncaging and resident lysosomal proteins were preferentially identified. Owing to the flexibility of our design featuring two photoreactions, this timing could readily be changed to longer intervals to allow for trafficking between organelles to occur in order to capture possible post-lysosome Sph transporters. We expect these tools to be broadly applicable not only to the study of sphingolipid export and related diseases and hope that this design sparks the generation of related tools to tackle questions regarding intraorganellar transfer of many different lipid species.

## Acknowledgements

The authors thank Wim Annaert (KU Leuven) for providing the HeLa NPC1^−/−^ cell line and Britta Brügger (Heidelberg University) for providing the HeLa SGPL1^−/−^ cell line. The work of J.A., J.N. and D.H. was funded by the Deutsche Forschungsgemeinschaft (DFG, German Research Foundation, project number 112927078 – TRR83) and the work of P.S. is supported by the Deutsche Forschungsgemeinschaft (DFG Research Group 2625; SA683/10-1).

## Author contributions

DH conceived and supervised the project. JA and JN synthesized the compounds. JN investigated protein-lipid interactions and prepared the samples for mass spectrometric analysis. JA performed thin-layer chromatography experiments together with JW as well as confocal microscopy experiments. PH performed LC-MS/MS measurements which were analysed by FS. SH generated LIMP2-KO cell lines with supervision from PS. JA and DH wrote the manuscript and all authors commented on the manuscript.

## Conflict of interest

The authors declare that they have no conflict of interest.

---

**Figure EV1.**
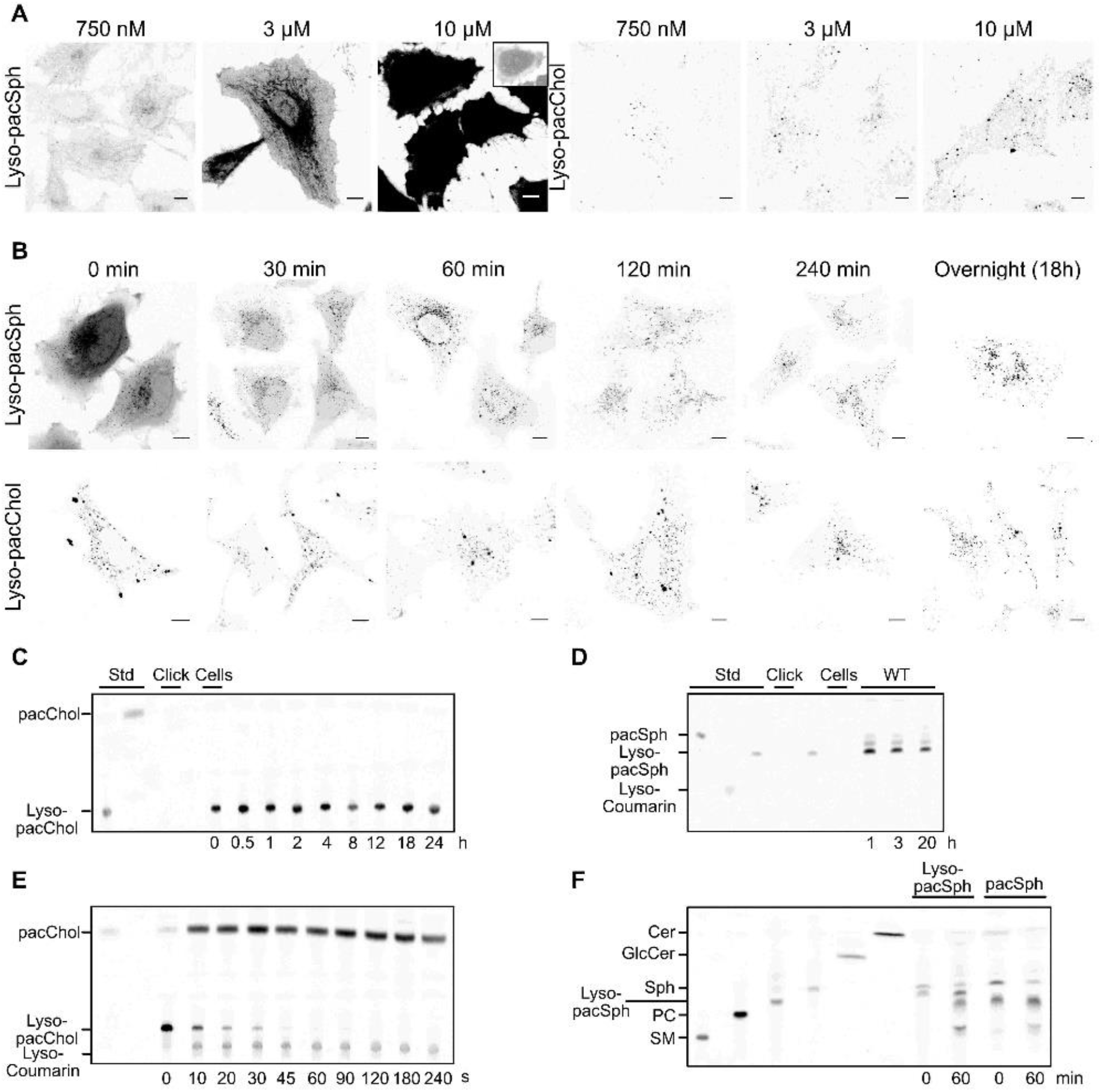
Characterisation of Lyso-pacSph and Lyso-pacChol. **A.** Concentration-dependent localisation of Lyso-pacSph and Lyso-pacChol. Live cell imaging of the coumarin fluorescence in HeLa cells pulsed with varying concentrations of Lyso-pacSph or Lyso-pacChol, chased for 30 minutes and imaged using the same microscopy settings. Due to image saturation at 10 μM Lyso-pacSph, the image was aquired again using lower laser power (small insert on the top right). **B.** Time-dependent localisation of Lyso-pacSph and Lyso-pacChol. Live cell imaging of the coumarin fluorescence in HeLa cells pulsed with Lyso-pacSph or Lyso-pacChol (10 μM) and chased and imaged for indicated times. **C, D.** Stability of Lyso-pacChol and Lyso-pacSph in cells. HeLa cells were labelled with Lyso-pacChol or Lyso-pacSph (10 μM) for 45 and 1h respectively, then chased for the indicated times. The probe was extracted, clicked with 3-azido-7-hydroxycoumarin and visualised by TLC. **E.** Uncaging efficiency of Lyso-pacChol in aqueous solution by TLC. Aqueous solutions of 10 μM Lyso-pacSph were irradiated with 405 nm light for increasing durations. Then, probe was extracted, clicked to 3-azido-7-hydroxycoumarin and visualised by TLC. **F.** Metabolite identification by TLC. HeLa WT cells were pulsed with 10 μM lyso-pacSph for 1h, chased overnight and uncaged. Lipids were extracted either directly upon uncaging (0 min) or after 60 min of incubation (60 min), clicked to 3-azido-7-hydroxycoumarin and visualised by TLC. To identify the lipid metabolites seen on the TLC, standards for ceramide (Cer), glucosylceramide (GlcCer), sphingosine (Sph), phosphatidylcholine (PC) and sphingomyelin (SM), clicked to 3-azido-7-hydroxycoumarin were spotted on the same plate.

**Figure EV2.**
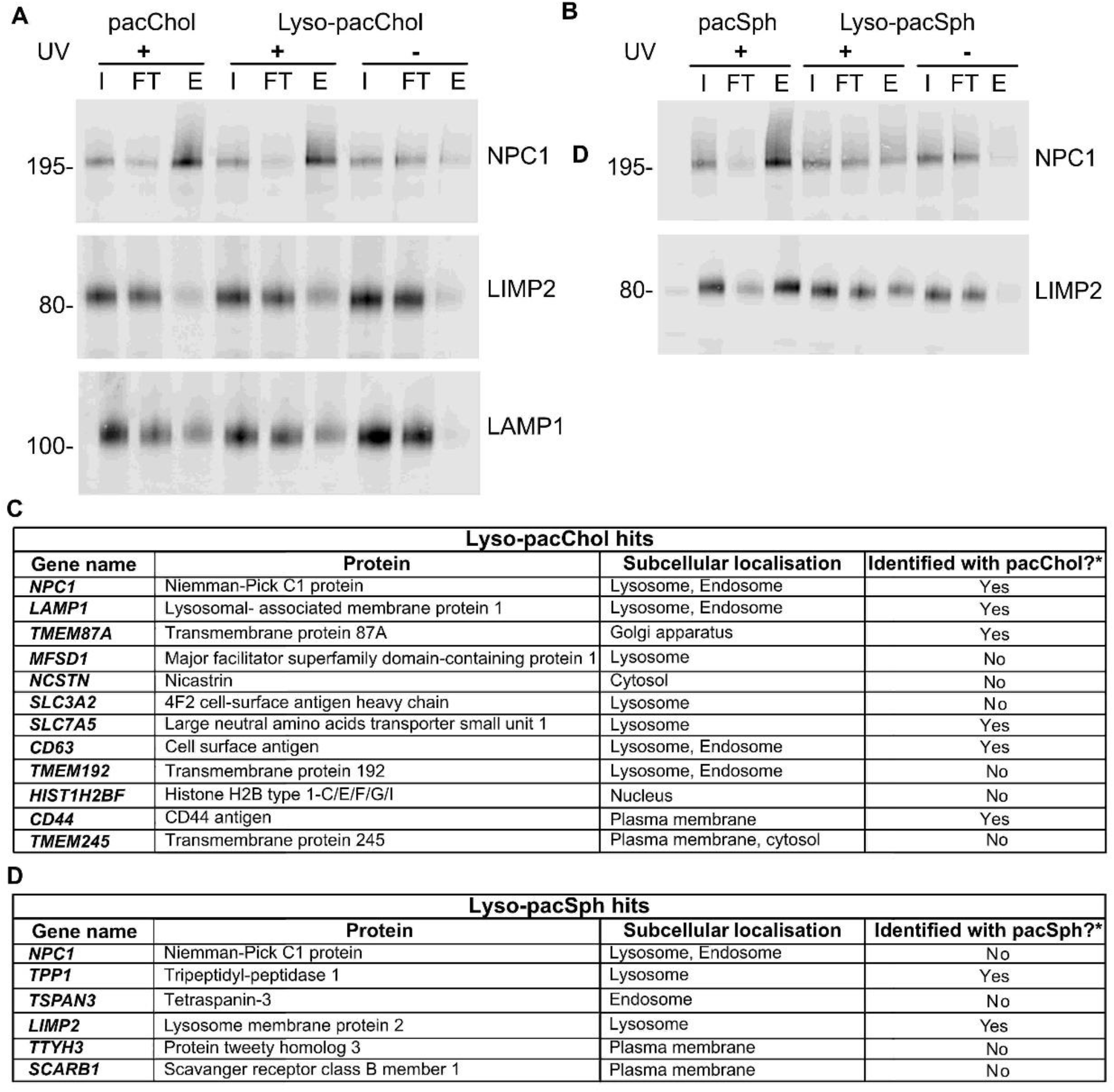
Application of lyso-pacChol and lyso-pacSph for identification of protein-lipid interactions. **A.** Identification of known cholesterol-binding proteins using lyso-pacChol and the globally distributed pacChol. Lysates of HeLa cells treated with pacChol and lyso-pacChol were uncaged, following by crosslinking (+UV) or no crosslinking (-UV) steps. Upon cell lysis and click reaction to biotin azide, the protein-lipid complexes were enriched by Streptavidin immunoprecipitation. The inputs (I, 10%), flow-throughs (FT, 10%) and eluates (E, 100%) were immunoblotted against NPC1, LIMP2, and LAMP1, respectively. **B.** Biochemical validation of novel lysosomal sphingosine interactors via Western blot. HeLa cells were treated with pacSph and Lyso-pacSph, respectively, crosslinked, lysed and subjected to click-reaction using biotin-azide. The biotinylated-protein lipid complexes were enriched by Streptavidin immunoprecipitation, the inputs (I), flow-throughs (FT) and eluates (E) of which were analysed by immunoblotting against NPC1 and LIMP2. **C. D.** List of identified hits, above threshold from the chemeoproteomic screen. Hits were analysed, anotated to their respective subcellular localisation and compared to previous proteome-wide mapping of pacSph and pacChol interacting proteins. *(Haberkant *et al.* 2016, Hulce *et al.* 2013)

**Figure EV3.**
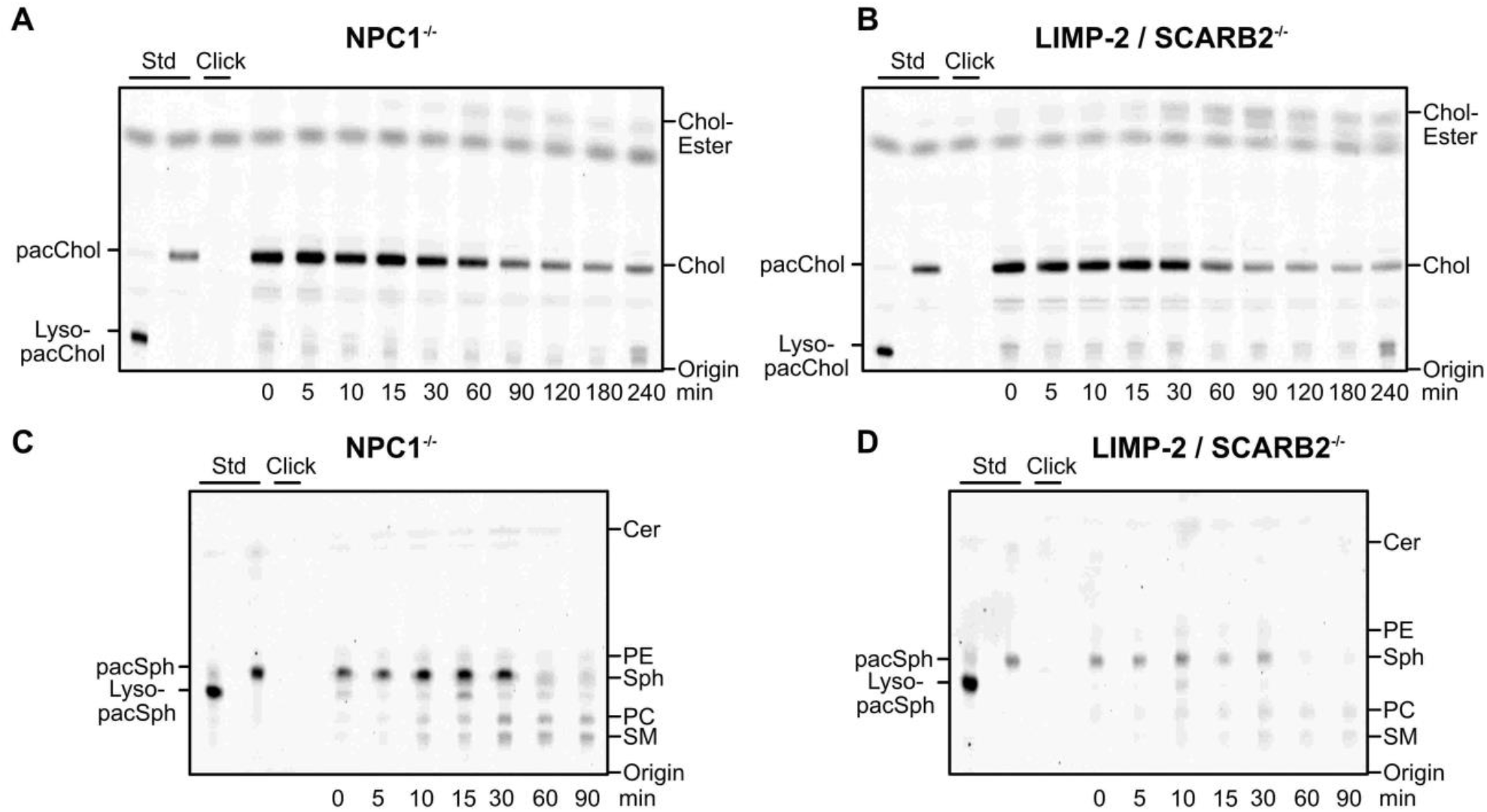
Lyso-pacChol and Lyso-pacSph metabolism in WT, NPC1 KO and LIMP2 KO cells by thin-layer chromatography. **A, B.** Post lysosomal metabolism of Lyso-pacChol. HeLa NPC1-deficient **(A)** and HeLa LIMP-2 deficient **(B)** cells were labelled with Lyso-pacChol (10 μM) for 45 min and chased overnight. Upon uncaging, cells were chased and lipids extracted at indicated times, clicked with 3-azido-7-hydroxycoumarin and visualised by thin-layer chromatography.**C,D.** Post lysosomal metabolism of Lyso-pacChol. HeLa NPC1-deficient **(C)** and HeLa LIMP-2 deficient **(D)** cells were labelled with Lyso-pacSph (10 μM) for 60 min and chased overnight. Upon uncaging, cells were chased and lipids extracted at indicated times, clicked with 3-azido-7-hydroxycoumarin and visualised by thin-layer chromatography.

**Figure EV4.1.**
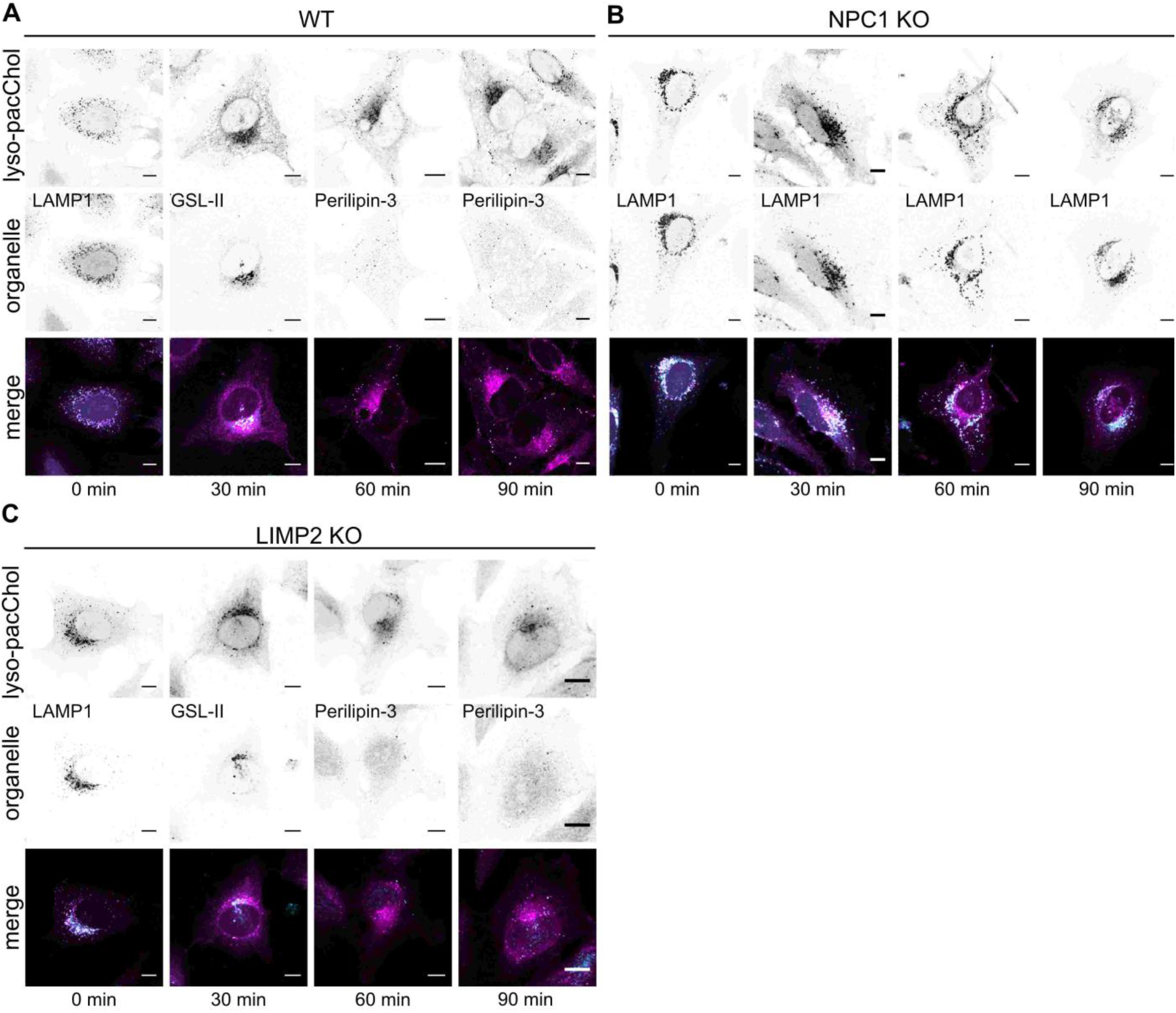
Visualisation of lysosomal cholesterol export across organelles. Identification of the subcellular localization of lyso-pacChol in WT **(A)**, NPC1 **(B)** and LIMP2 **(C)**-deficient HeLa cells. Confocal images of cells labeled with Lyso-pacChol (10 μM) for 45 min and chased overnight. Upon uncaging, cells were chased for the indicated times, crosslinked, 4% PFA/ 0.1% glyoxal fixed, functionalised with AlexaFluor 488-Picolyl-Azide and subjected to immunofluorescence staining with antibody against LAMP1 (lysosomes), Periliplin-3 (lipid droplets) or Lectin GS-II From *Griffonia simplicifolia*, Alexa Fluor-647 (Golgi). Scale bar: 10 μm

**Figure EV4.2.**
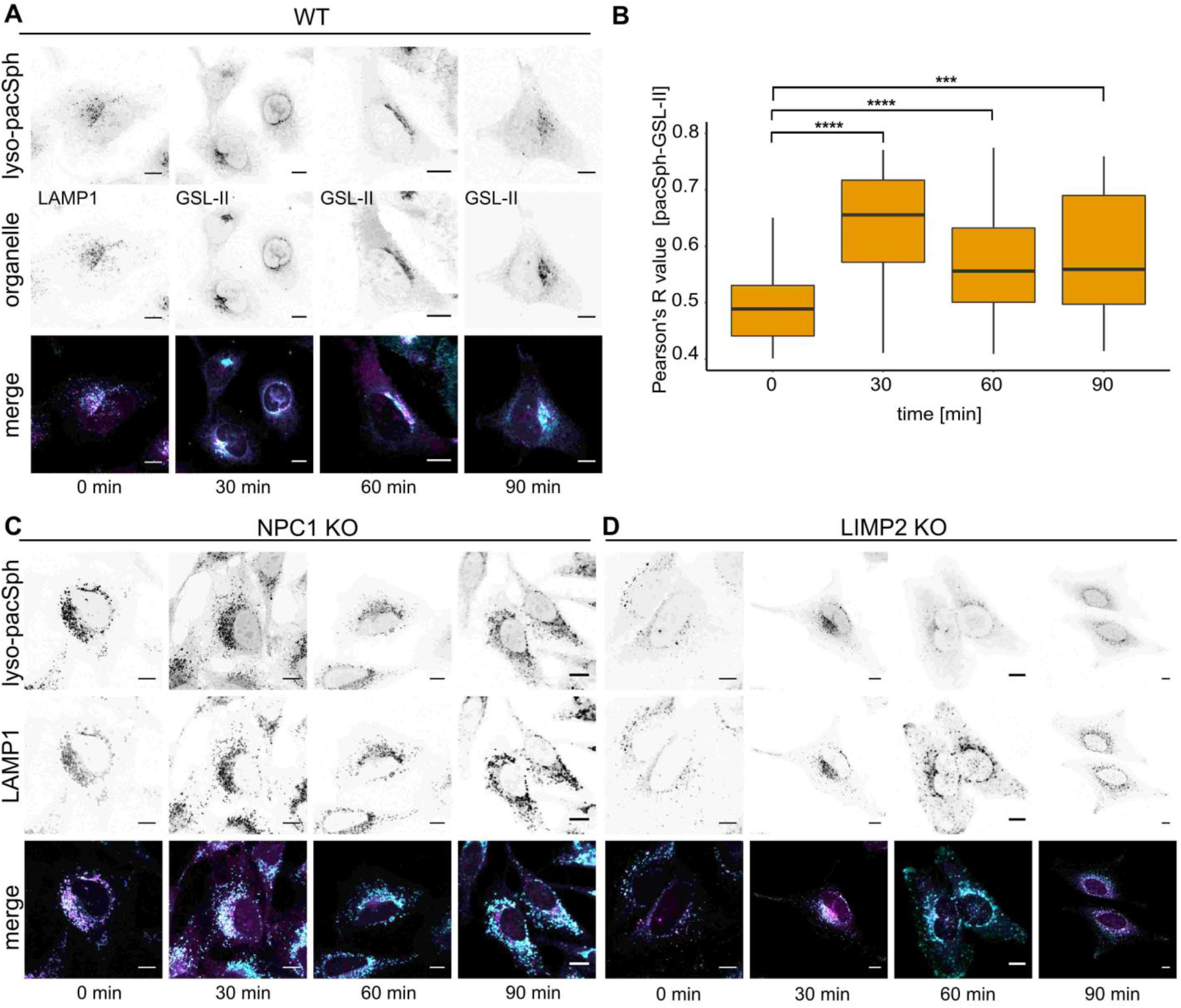
Visualisation of lysosomal sphingosine export across organelles. Identification of the subcellular localization of lyso-pacSph in WT **(A)**, NPC1 **(B)** and LIMP2 **(C)**-deficient HeLa cells. Confocal images of cells labeled with lyso-pacSph (10 μM) for 1h and chased overnight. Upon uncaging, cells were chased for the indicated times, crosslinked, methanol fixed, functionalised with AlexaFluor 594-Picolyl-Azide and subjected to immunofluorescence staining with antibody against LAMP1 (lysosomes) and Lectin GS-II From *Griffonia simplicifolia*, Alexa Fluor647 (Golgi). Scale bar: 10 μm **B.** Quantification of sphingosine-Golgi colocalisation. Pearson’s R value of non-thresholded images from pacSph vs Lectin (GS-II) immunofluorescence, calculated for each timepoint (n ≥ 44) using the Coloc 2 feature from Fiji. (ns P-value > 0.5 *P ≤ 0.05 **P-value ≤ 0.01 ***P-value ≤ 0.001 ****P-value ≤ 0.0001)

**Figure EV5.1.**
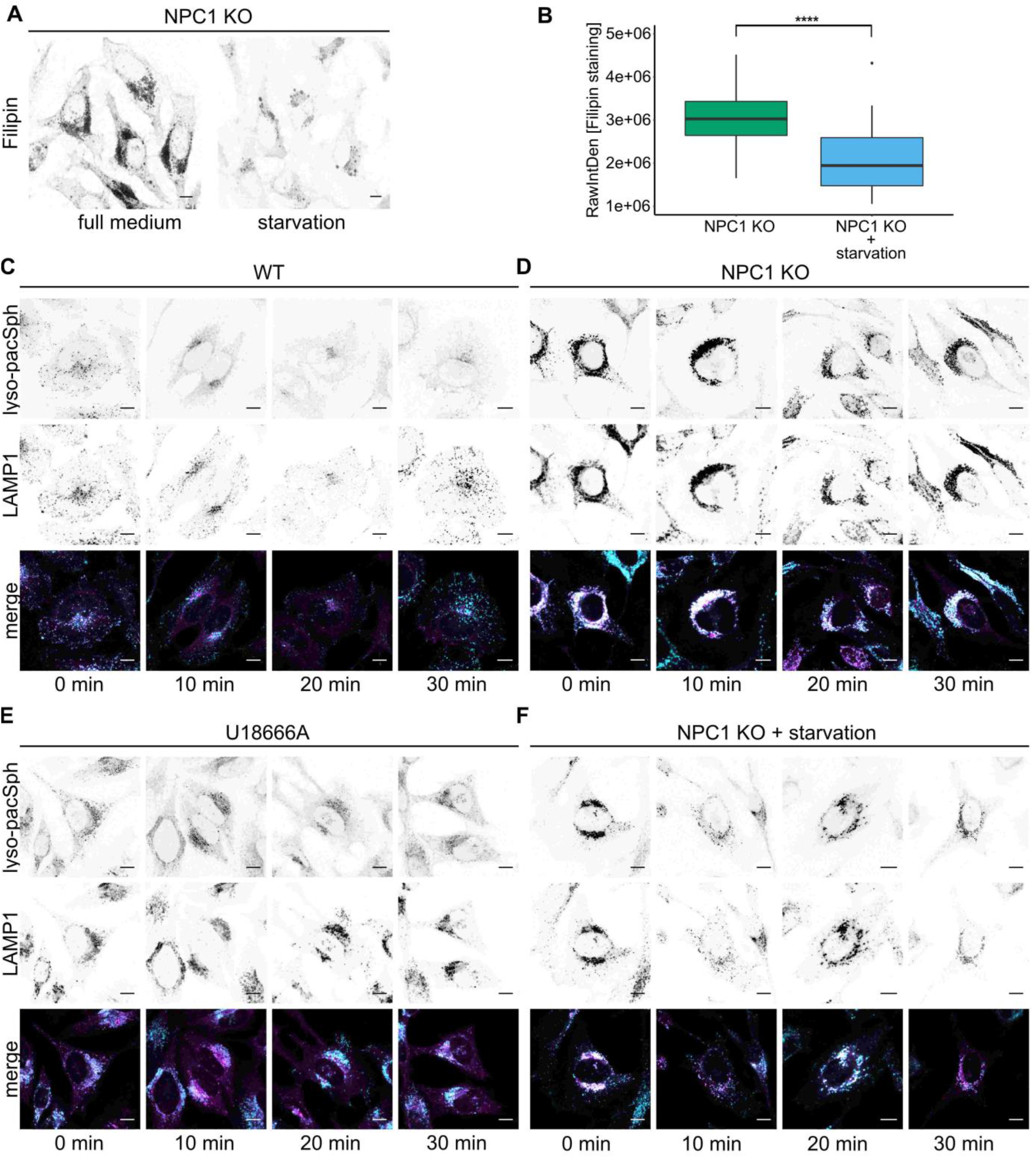
Sphingosine accumulation is a direct effect from loss of NPC1. Effect of lipoprotein deficient medium. Confocal images of HeLa NPC1 KO cells **(A)** in regular medium conditions or incubated with lipoprotein deficient medium 48h prior imaging. Quantification of cholesterol levels via filipin staining **(B)**. Raw intensity density (RawIntDen) of non-thresholded images from filipin staining signal, calculated for each condition (n ≥ 34) using a ROI in Fiji. (****P-value ≤ 0.0001) Confocal images of Lyso-pacSph in cellular models of NPC1. HeLa WT cells **(C)**, HeLa NPC1 KO cells **(D)**, HeLa WT cells treated with 0.5 μg/mL U18666A 24h prior to imaging (U18666A) **(E)** and HeLa NPC1 KO cells incubated with lipoprotein deficient medium 48h prior to imaging (NPC1 KO + starvation) **(F)** were labeled with lyso-pacSph (1.25 μM) for 1h and chased overnight. Upon uncaging, chase-crosslinking experiments were performed from 0 to 30 min. Cells were fixed with methanol, functionalised with AlexaFluor 594-Picolyl-Azide and subjected to immunofluorescence staining with antibody against LAMP1 (lysosomes). Scale bar: 10 μm

**Figure EV5.2.**
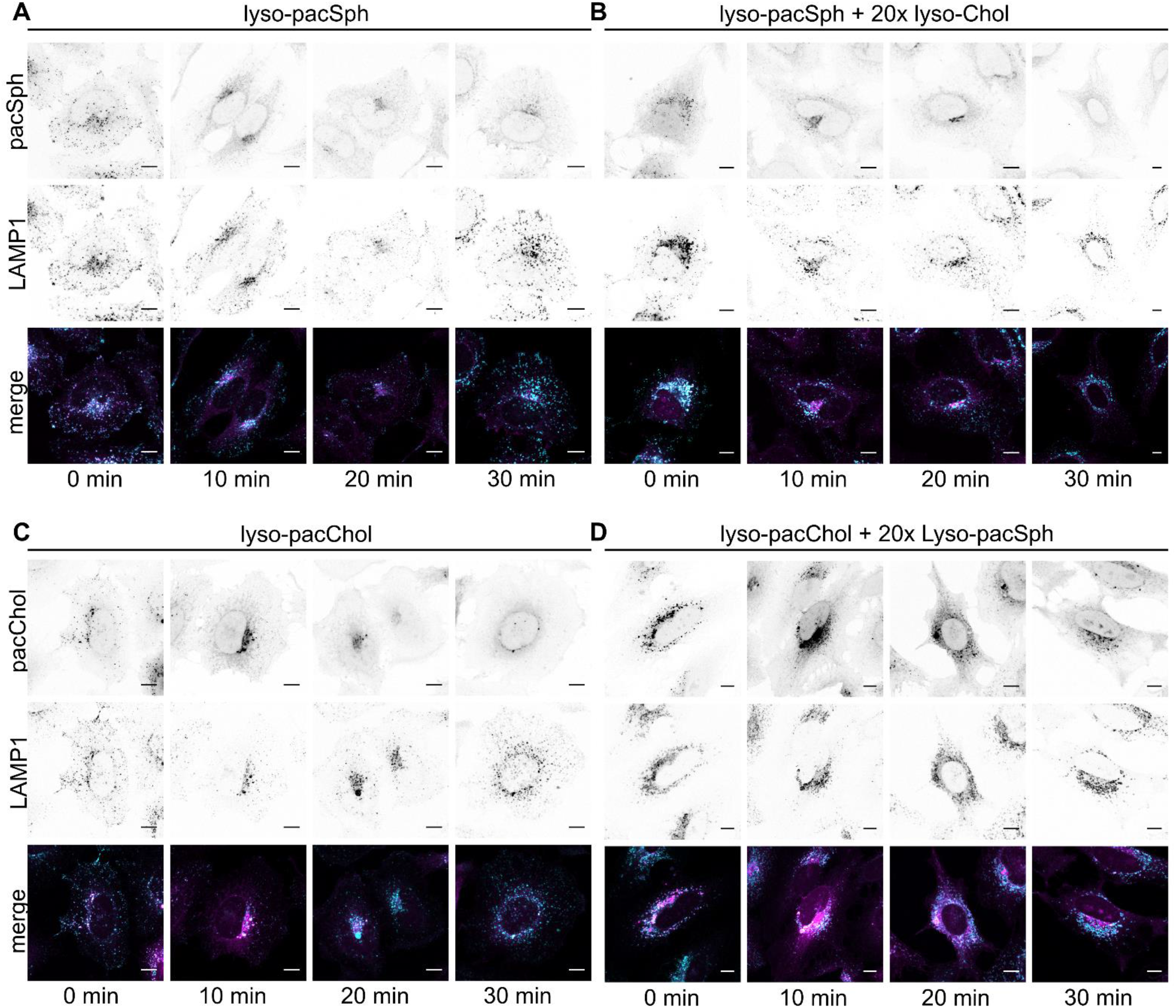
Lysosomal sphingosine accumulation can re-create an NPC1-like scenario. Confocal images of Lyso-pacSph and Lyso-pacChol. HeLa WT cells were labeled with Lyso-pacSph (1.25 μM) **(A)**, Lyso-pacSph (1.25 μM) and Lyso-Chol (25 μM) **(B)**, Lyso-pacChol (750 nM) **(C)** and Lyso-pacChol (750 nM) and Lyso-Sph (15 μM) **(D)** for 1h and chased overnight. Upon uncaging, chase-crosslinking experiments were performed from 0 to 30 min. Cells were fixed, functionalised with AlexaFluor 594-Picolyl-Azide **(A, B)** or AlexaFluor 488-Picolyl-Azide **(C,D)** and stained with an antibody against LAMP1 (lysosomes). Scale bar: 10 μm

## References

Antonny B, Bigay J & Mesmin B (2018) The Oxysterol-Binding Protein Cycle: Burning off PI(4)P to Transport Cholesterol. Annual Review of Biochemistry 87: 809–837

Ashcroft RG, Thulborn KR, Smith JR, Coster HGL & Sawyer WH (1980) Perturbations to lipid bilayers by spectroscopic probes as determined by dielectric measurements. Biochimica et Biophysica Acta - Biomembranes 602: 299–308

Ballabio A & Bonifacino JS (2020) Lysosomes as dynamic regulators of cell and organismal homeostasis. Nature Reviews Molecular Cell Biology 21: 101–118 doi:10.1038/s41580-019-0185-4 [PREPRINT]

Bektas M, Laura Allende M, Lee BG, Chen WP, Amar MJ, Remaley AT, Saba JD & Proia RL (2010) Sphingosine 1-phosphate lyase deficiency disrupts lipid homeostasis in liver. Journal of Biological Chemistry 285: 10880–10889

Blom T, Li Z, Bittman R, Somerharju P & Ikonen E (2012) Tracking Sphingosine Metabolism and Transport in Sphingolipidoses: NPC1 Deficiency as a Test Case. Traffic 13: 1234–1243

Bockelmann S, Mina JGM, Korneev S, Hassan DG, Müller D, Hilderink A, Vlieg HC, Raijmakers R, Heck AJR, Per H, et al (2018) A search for ceramide binding proteins using bifunctional lipid analogs yields CERT-related protein StarD7. Journal of Lipid Research 59: 515–530

Brown MS & Goldstein JL (1986) A receptor-mediated pathway for cholesterol homeostasis. Science 232: 34–47

Carstea ED (1996) Niemann-Pick C1 Disease Gene: Homology to Mediators of Cholesterol Homeostasis. Science 370: 1

Chattopadhyay A (1990) Chemistry and biology of N-(7-nitrobenz-2-oxa-1,3-diazol-4-yl)-labeled lipids: fluorescent probes of biological and model membranes. Chemistry and Physics of Lipids 53: 1–15

Conrad KS, Cheng TW, Ysselstein D, Heybrock S, Hoth LR, Chrunyk BA, Am Ende CW, Krainc D, Schwake M, Saftig P, et al (2017) Lysosomal integral membrane protein-2 as a phospholipid receptor revealed by biophysical and cellular studies. Nature Communications 8

Du X, Kumar J, Ferguson C, Schulz TA, Ong YS, Hong W, Prinz WA, Parton RG, Brown AJ & Yang H (2011) A role for oxysterol-binding protein-related protein 5 in endosomal cholesterol trafficking. Journal of Cell Biology 192: 121–135

Feng S, Harayama T, Chang D, Hannich JT, Winssinger N & Riezman H (2019a) Lysosome-targeted photoactivation reveals local sphingosine metabolism signatures. Chemical Science 10: 2253–2258

Feng S, Harayama T, Chang D, Hannich JT, Winssinger N & Riezman H (2019b) Lysosome-targeted photoactivation reveals local sphingosine metabolism signatures. Chemical Science 10: 2253–2258

Futerman AH & Riezman H (2005) The ins and outs of sphingolipid synthesis. Trends in Cell Biology 15: 312–318

Gerl MJ, Bittl V, Kirchner S, Sachsenheimer T, Brunner HL, Lüchtenborg C, Özbalci C, Wiedemann H, Wegehingel S, Nickel W, et al (2016) Sphingosine-1-phosphate lyase deficient cells as a tool to study protein lipid interactions. PLoS ONE 11

Gillard BK, Clement RG & Marcus DM (1998) Variations among cell lines in the synthesis of sphingolipids in de novo and recycling pathways. Glycobiology 8: 885–890

Girik V, Feng S, Hariri H, Henne M & Riezman H (2021) Vacuole-specific lipid release for tracking intracellular lipid metabo-lism and transport in Saccharomyces cerevisiae. bioRxiv: 2021.05.04.442581

Goldstein JL, DeBose-Boyd RA & Brown MS (2006) Protein sensors for membrane sterols. Cell 124: 35–46

Haberkant P, Raijmakers R, Wildwater M, Sachsenheimer T, Brügger B, Maeda K, Houweling M, Gavin AC, Schultz C, van Meer G, et al (2013) In vivo profiling and visualization of cellular protein-lipid interactions using bifunctional fatty acids. Angewandte Chemie - International Edition 52: 4033–4038

Haberkant P, Stein F, Höglinger D, Gerl MJ, Brügger B, van Veldhoven PP, Krijgsveld J, Gavin AC & Schultz C (2016) Bifunctional Sphingosine for Cell-Based Analysis of Protein-Sphingolipid Interactions. ACS Chemical Biology 11: 222–230

Hannun YA & Bell RM (1989) Functions of sphingolipids and sphingolipid breakdown products in cellular regulation. Science 243: 500–507

Hannun YA & Obeid LM (2008) Principles of bioactive lipid signalling: Lessons from sphingolipids. Nature Reviews Molecular Cell Biology 9: 139–150

Heybrock S, Kanerva K, Meng Y, Ing C, Liang A, Xiong ZJ, Weng X, Ah Kim Y, Collins R, Trimble W, et al (2019) Lysosomal integral membrane protein-2 (LIMP-2/SCARB2) is involved in lysosomal cholesterol export. Nature Communications 10

Höglinger D, Burgoyne T, Sanchez-Heras E, Hartwig P, Colaco A, Newton J, Futter CE, Spiegel S, Platt FM & Eden ER (2019) NPC1 regulates ER contacts with endocytic organelles to mediate cholesterol egress. Nature Communications 10: 1–14

Höglinger D, Nadler A, Haberkant P, Kirkpatrick J, Schifferer M, Stein F, Hauke S, Porter FD & Schultz C (2017) Trifunctional lipid probes for comprehensive studies of single lipid species in living cells. Proceedings of the National Academy of Sciences of the United States of America 114: 1566–1571

Höglinger D, Nadler A & Schultz C (2014) Caged lipids as tools for investigating cellular signaling. Biochimica et Biophysica Acta - Molecular and Cell Biology of Lipids 1841: 1085–1096

Hölttä-Vuori M, Uronen RL, Repakova J, Salonen E, Vattulainen I, Panula P, Li Z, Bittman R & Ikonen E (2008) BODIPY-cholesterol: A new tool to visualize sterol trafficking in living cells and organisms. Traffic 9: 1839–1849

Hulce JJ, Cognetta AB, Niphakis MJ, Tully SE & Cravatt BF (2013) Proteome-wide mapping of cholesterol-interacting proteins in mammalian cells. Nature Methods 10: 259–264

Infante RE, Wang ML, Radhakrishnan A, Hyock JK, Brown MS & Goldstein JL (2008) NPC2 facilitates bidirectional transfer of cholesterol between NPC1 and lipid bilayers, a step in cholesterol egress from lysosomes. Proceedings of the National Academy of Sciences of the United States of America 105: 15287–15292

Kaufmann AM & Krise JP (2007) Lysosomal sequestration of amine-containing drugs: Analysis and therapeutic implications. Journal of Pharmaceutical Sciences 96: 729–746 doi:10.1002/jps.20792 [PREPRINT]

Kitatani K, Idkowiak-Baldys J & Hannun YA (2008a) The sphingolipid salvage pathway in ceramide metabolism and signaling. Cellular Signalling 20: 1010–1018

Kitatani K, Idkowiak-Baldys J & Hannun YA (2008b) The sphingolipid salvage pathway in ceramide metabolism and signaling. Cellular Signalling 20: 1010–1018 doi:10.1016/j.cellsig.2007.12.006 [PREPRINT]

Kwon HJ, Abi-Mosleh L, Wang ML, Deisenhofer J, Goldstein JL, Brown MS & Infante RE (2009) Structure of N-Terminal Domain of NPC1 Reveals Distinct Subdomains for Binding and Transfer of Cholesterol. Cell 137: 1213–1224

Li J & Pfeffer SR (2016) Lysosomal membrane glycoproteins bind cholesterol and contribute to lysosomal cholesterol export. eLife

Lloyd-Evans E, Morgan AJ, He X, Smith DA, Elliot-Smith E, Sillence DJ, Churchill GC, Schuchman EH, Galione A & Platt FM (2008) Niemann-Pick disease type C1 is a sphingosine storage disease that causes deregulation of lysosomal calcium. Nature Medicine 14: 1247–1255

Lu F, Liang Q, Abi-Mosleh L, Das A, de Brabander JK, Goldstein JL & Brown MS (2015) Identification of NPC1 as the target of U18666A, an inhibitor of lysosomal cholesterol export and Ebola infection. eLife 4: 1–16

Maier O, Oberle V & Hoekstra D (2002) Fluorescent lipid probes: some properties and applications (a review)

Marques ARA & Saftig P (2019) Lysosomal storage disorders – challenges, concepts and avenues for therapy: Beyond rare diseases. Journal of Cell Science 132

Meng Y, Heybrock S, Neculai D & Saftig P (2020) Cholesterol Handling in Lysosomes and Beyond. Trends in Cell Biology 30: 452–466 doi:10.1016/j.tcb.2020.02.007 [PREPRINT]

Nadler A, Yushchenko DA, Müller R, Stein F, Feng S, Mulle C, Carta M & Schultz C (2015) Exclusive photorelease of signalling lipids at the plasma membrane. Nature Communications 6: 1–10

Neculai D, Schwake M, Ravichandran M, Zunke F, Collins RF, Peters J, Neculai M, Plumb J, Loppnau P, Pizarro JC, et al (2013) Structure of LIMP-2 provides functional insights with implications for SR-BI and CD36. Nature 504: 172–176

Nguyen Trinh M, Brown MS, Seemann J, Goldstein JL & Lu F (2018) Lysosomal cholesterol export reconstituted from fragments of Niemann-Pick C1. eLife 7: 1–14

Pagano RE, Watanabe R, Wheatley C & Dominguez M (2000) Applications of BIODIPY-sphingolipid analogs to study lipid traffic and metabolism in cells Elsevier Masson SAS

Platt FM (2018) Emptying the stores: Lysosomal diseases and therapeutic strategies. Nature Reviews Drug Discovery 17: 133–150

Rodriguez-Lafrasse C, Rousson R, Pentchev PG, Louisot P & Vanier MT (1994) Free sphingoid bases in tissues from patients with type C Niemann-Pick disease and other lysosomal storage disorders. BBA - Molecular Basis of Disease 1226: 138–144

Saha P, Shumate JL, Caldwell JG, Elghobashi-Meinhardt N, Lu A, Zhang L, Olsson NE, Elias JE & Pfeffer SR (2020) Inter-domain dynamics drive cholesterol transport by NPC1 and NPC1L1 proteins. bioRxiv 3: 1–28

Saheki Y, Bian X, Schauder CM, Sawaki Y, Surma MA, Klose C, Pincet F, Reinisch KM & De Camilli P (2016) Control of plasma membrane lipid homeostasis by the extended synaptotagmins. Nature Cell Biology 18: 504–515

Schoop V, Martello A, Eden ER & Höglinger D (2021) Cellular cholesterol and how to find it. Biochimica et Biophysica Acta - Molecular and Cell Biology of Lipids 1866 doi:10.1016/j.bbalip.2021.158989 [PREPRINT]

Schwarzmann G, Arenz C & Sandhoff K (2014) Labeled chemical biology tools for investigating sphingolipid metabolism, trafficking and interaction with lipids and proteins. Biochimica et Biophysica Acta - Molecular and Cell Biology of Lipids 1841: 1161–1173

Tharkeshwar AK, Trekker J, Vermeire W, Pauwels J, Sannerud R, Priestman DA, Te Vruchte D, Vints K, Baatsen P, Decuypere JP, et al (2017) A novel approach to analyze lysosomal dysfunctions through subcellular proteomics and lipidomics: The case of NPC1 deficiency. Scientific Reports 7: 1–20

Thiele C & Spandl J (2008) Cell biology of lipid droplets. Current Opinion in Cell Biology 20: 378–385

Vanharanta L, Peränen J, Pfisterer SG, Enkavi G, Vattulainen I & Ikonen E (2020) High-content imaging and structure-based predictions reveal functional differences between Niemann-Pick C1 variants. Traffic 21: 386–397

Wagner N, Stephan M, Höglinger D & Nadler A (2018) A Click Cage: Organelle-Specific Uncaging of Lipid Messengers. Angewandte Chemie - International Edition 57: 13339–13343

Winkler MBL, Kidmose RT, Szomek M, Thaysen K, Rawson S, Muench SP, Wüstner D & Pedersen BP (2019) Structural Insight into Eukaryotic Sterol Transport through Niemann-Pick Type C Proteins. Cell 179: 485–497.e18

Xu S, Benoff B, Liou HL, Lobel P & Stock AM (2007) Structural basis of sterol binding by NPC2, a lysosomal protein deficient in Niemann-Pick type C2 disease. Journal of Biological Chemistry 282: 23525–23531

Xu W, Zeng Z, Jiang JH, Chang YT & Yuan L (2016) Discerning the Chemistry in Individual Organelles with Small-Molecule Fluorescent Probes. Angewandte Chemie - International Edition 55: 13658–13699 doi:10.1002/anie.201510721 [PREPRINT]

Zhang X, Zheng N & Rosania GR (2008) Simulation-based cheminformatic analysis of organelle-targeted molecules: Lysosomotropic monobasic amines. Journal of Computer-Aided Molecular Design 22: 629–645

